# Overcoming a “forbidden phenotype”: The parrot’s head supports, propels, and powers tripedal locomotion

**DOI:** 10.1101/2021.12.22.473737

**Authors:** Melody W. Young, Edwin Dickinson, Nicholas D. Flaim, Michael C. Granatosky

## Abstract

No vertebrate, living or extinct, is known to have possessed an odd number of limbs. Despite this “forbidden phenotype", gaits that utilize odd numbers of limbs (e.g., tripedalism or pentapedalism) have evolved in both avian and mammalian lineages. Tripedal locomotion is commonly employed by parrots during climbing, who utilize their beaks as an additional support. However, it is unclear whether the beak functions simply as a stabilizing hook, or as a propulsive limb. Here, we present data on kinetics of tripedal climbing in six rosy-faced lovebirds (*Agapornis rosiecollis*). Our findings demonstrate that parrots utilize cyclical tripedal gaits when climbing and the beak and hindlimbs generate comparable propulsive and tangential substrate reaction forces and power. Propulsive and tangential forces generated by the beak are of equal or greater magnitudes to those forces generated by the forelimbs of humans and non-human primates during vertical climbing. We conclude that the feeding apparatus and neck musculature of parrots has been co-opted to function biomechanically as a third limb during vertical climbing. We hypothesize that this exaptation required substantive alterations to the neuromuscular system including enhanced force-generating capabilities of the neck musculature and modifications to limb central pattern generators.

## Introduction

There is no odd-legged vertebrate alive today, nor is there any evidence of such a creature ever existing in the fossil record (1). This “forbidden phenotype” is inherently reliant on the presence of a midline limb, but phylogenetic constraints of bilateral limb development circumvent such a possibility (1). Despite the lack of odd-limbed tetrapods, tripedal and pentapedal gaits have emerged in multiple lineages, including mammals and birds (1–3).

One well-known, but anecdotal (but see 2), example of tripedal locomotion is the climbing behaviors of parrots (Order: Psittaciformes) (1, 4). Unable to use their wings as grasping forelimbs, parrots have evolved to be resourceful climbers by co-opting their feeding apparatus as an additional “limb” (2). However, it is unclear if parrots use their beaks to simply hook into the substrate (serving a primarily stabilizing function), or whether the head function is a true evolutionary novelty, exapting the head as a functional limb to propel and power the body in a cyclical manner. Here we present kinetic data to quantitatively assess the functional role of the head during the tripedal climbing gaits of parrots.

## Results

When climbing, parrots exhibit cyclical tripedal gaits (Figure 1), utilizing the beak as an effective third limb. After accounting for speed, the beak (70.6 ± 20.2 %bw; *n* = 123) generated a propulsive force no different than the hindlimb(s) (74.4 ± 20.4 %bw; *n* = 111; *p*=0.357; Figure 2). While the beak and hindlimbs exerted exclusively propulsive forces, the tail served a braking function (−14.7 ± 7.1 %bw; *n* = 98). The beak also experiences tangential forces (46.9 ± 14.5 %bw) similar in magnitude to the hindlimb(s) (45.4 ± 15.2 %bw, *p*=0.593). However, while the beak’s tangential forces are purely tensile in nature, the hindlimb transitions from tensile to compressive loading depending on the foot’s placement during the stride (5). Although contributing a lesser degree than the beak and hindlimbs, the tail provided additional compressive tangential forces (18.8 ± 6.83 %bw; *p* ≤ 0.001).

**Figure 1.**
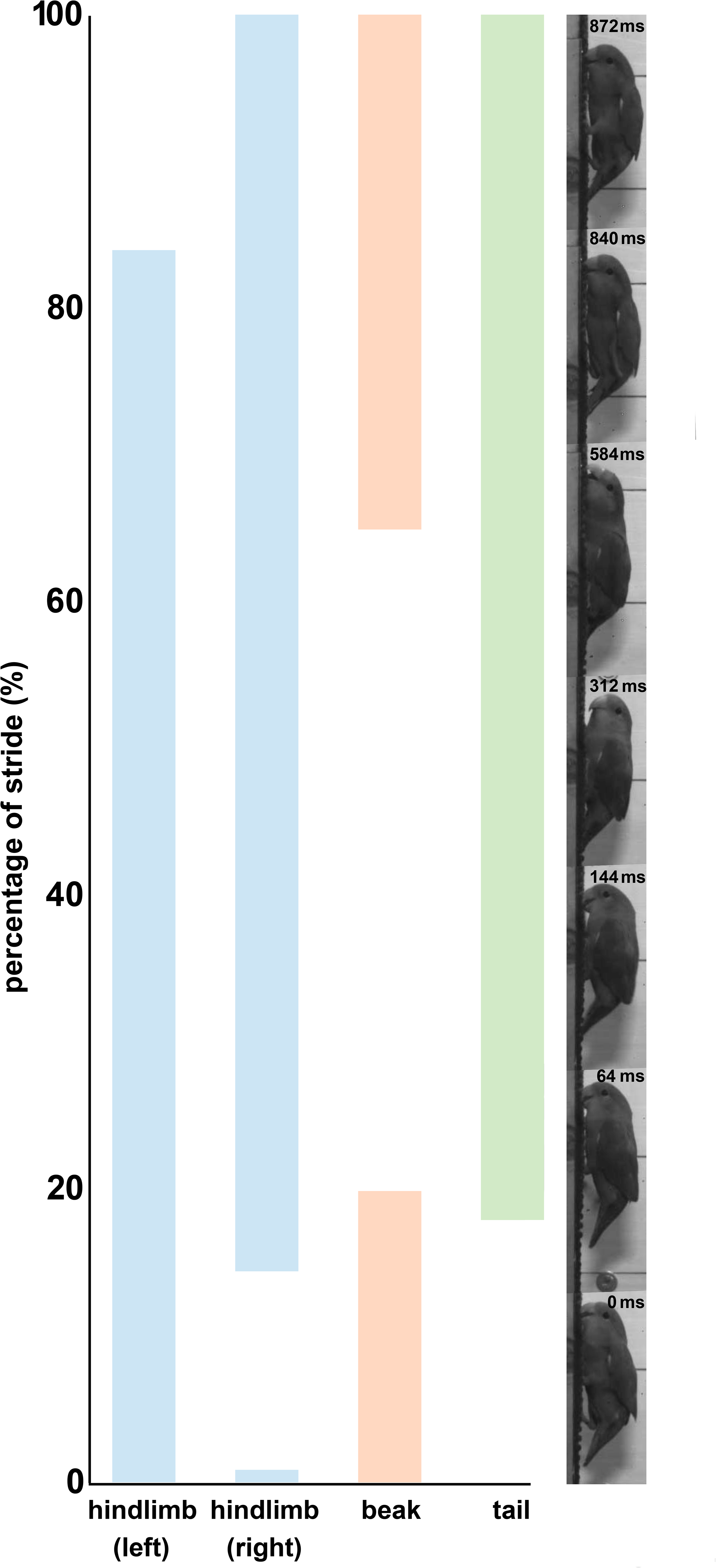
Representative cycle observed during vertical climbing in the rosy-faced lovebird (*Agapornis roseicollis*).

**Figure 2.**
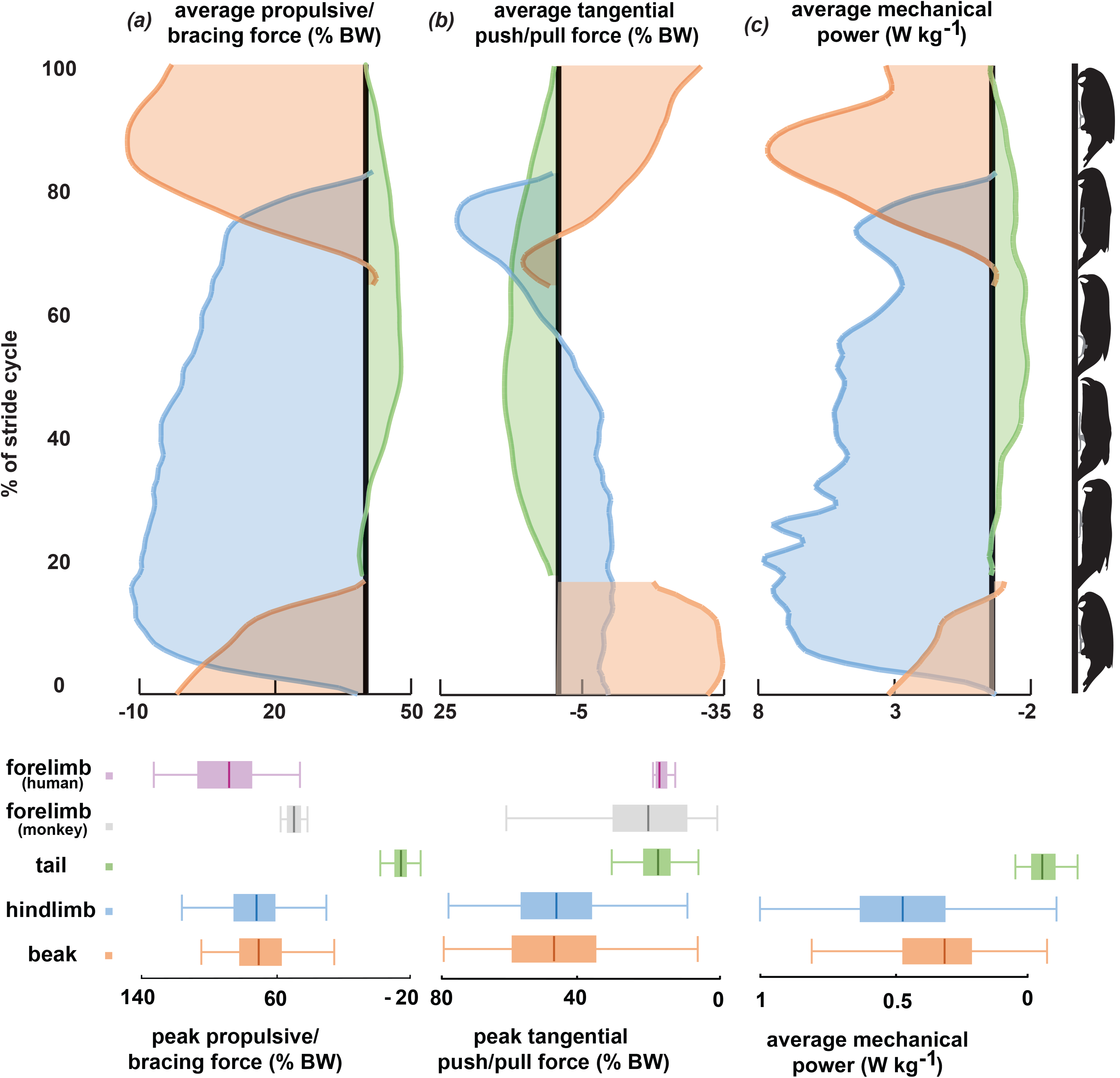
Force and power generation by the beak, hindlimb (left), and tail during vertical climbing in the rosy-faced lovebird (*Agapornis roseicollis*). The silhouettes placed on the right correspond with distinct events during a stride. (a) Average (top) and peak (bottom; box and whisker plots) propulsive/braking substrate reaction force (SRF) demonstrate that the beak (orange) provides equal propulsive force to the hindlimb (blue) and is comparable to human (1) and primate (2) forelimbs during climbing. (b) Tangential SRF illustrates that the beak provides an equal but opposite pulling force (top) to the pushing force of the hindlimb to balance on the vertical substrate. For comparative purposes, the absolute value of the beak tangential forces were used for all statistical analyses and displayed graphically. Peak tangential forces (bottom) of the beak and hindlimbs are greater than those produced by primates and humans during climbing. (c) Average single limb mechanical power demonstrates the beak performs mechanical work nearly equal to that of the hindlimb.

The beak generated a substantial magnitude of positive mechanical work (0.3 ± 0.2 W kg^−1^), almost matching that of the hindlimbs (0.5 ± 0.2 W kg^−1^). The tail, meanwhile, contributed only negative mechanical work (−0.1 W kg^−1^) All post-hoc comparisons *p* ≤ 0.001. These parrots are thus reliant on the combined contribution of positive power from the hindlimbs and beak to ascend vertical surfaces.

## Discussion

The feeding apparatus and neck musculature of parrots has been co-opted to function biomechanically as a third limb, conferring both the stability and propulsive force necessary to power a tripedal gait. In the tangential plane, the beak also serves to counteract the gravitational torque that threatens to pitch the animal away from the support, in a similar manner to the hindlimbs. Both weight adjusted propulsive and tangential forces in the head match, or even exceed, peak magnitudes in the forelimb during vertical climbing in primates (5) and rock climbing in humans (e.g., 8). By contrast, the tail of parrots contributes compressive bracing forces that result in negative mechanical work throughout the stride. As such, it is inappropriate to refer to the tail of climbing parrots as a limb. Instead, the tail functions as a simple stabilizing strut, bracing the body in a manner similar to descriptions of tail usage in climbing woodpeckers (7) and treecreepers (8), but not contributing to forward propulsion or power during locomotion.

Utilizing the head as a limb represents an evolutionary novelty that, to our knowledge, has only ever arisen within Psittaciformes. The evolution of such a behavioral innovation requires the exaptation of existing features of the feeding apparatus for climbing. To prevent pitching backwards off vertical substrates, parrots use their beaks to grasp onto the support. The force-generating capabilities of the parrot’s beak are well-known, and while not measured *in vivo*, estimates using muscle cross-sectional proportions suggest they can produce forces ~14x their body mass (9). This is more than sufficient to support the ~47%bw tangential tensile forces experienced during climbing. However, as a consequence of the stiffened thoracic and lumbar region in birds (10), propulsion must be completely driven by contraction of the neck musculature. Little is known about the force generating capabilities of neck musculature across vertebrates, but limited data collected from humans suggest maximal isometric neck extension force to be ~34%bw and neck flexion force ~19%bw (11). These data suggest that to provide the recorded ~71%bw of propulsive force recorded at the beak during climbing, parrots must have greatly enhanced the relative torque-generating capabilities of the neck musculature to facilitate the demands of tripedal locomotion.

Beyond the anatomical changes necessary to use the head as a third limb, considerable neuromuscular modifications must have evolved in parallel to permit such a radical innovation to locomotor patterning. While many birds climb vertical supports, only the parrots use their head as a third limb; by contrast, woodpeckers, nuthatches, and treecreepers ascend via synchronous hindlimb movements that essentially allow the animal to hop up the tree. Parrots rarely hop (4, 12), and appear to have lost this ability early their evolutionary history (12). The use of the beak as a third limb may be either a consequence of, or the causal factor driving, the loss of this ability.

Incorporating the head into the locomotor system constitutes a complication to the bilateral cycling of limb central pattern generators (CPGs) and raises the possibility of interlimb interference throughout the stride (13). However, it is possible that birds may not require radical neuromuscular innovation to incorporate movements of the head within the locomotor cycle due to the presence of head-bobbing [i.e., a conspicuous behavior in which the fore-aft movement of the head is coordinated with footfall during locomotion (14)]. Thus, parrots may simply have repurposed this neuromuscular to utilize the head as a third limb during climbing.

## Conclusion

Despite the absence of odd-limbed tetrapods, tripedal and/or pentapedal gaits have evolved in both avian and mammalian lineages. In this study, we present a kinetic analysis of tripedal locomotion in a climbing parrot. We demonstrate that, when climbing, these birds utilize cyclical tripedal gaits in which the beak functions as an effective third limb, generating comparable tangential and propulsive forces and power to the hindlimbs. We conclude that the feeding apparatus of parrots has been co-opted to function biomechanically as a third limb during vertical climbing. Utilizing the head as a limb represents an evolutionary novelty to Psittaciformes and may be facilitated by the neuromuscular modifications to limb CPGs and/or existing coupling between head and limb movements in birds.

## Materials and Methods

Beak, hindlimb, and tail fore-aft and tangential substrate reaction forces (SRFs) were collected from six rosy-faced lovebirds (*Agapornis roseicollis*) as the animals climbed an instrumented (HE6X6; Advanced Mechanical Technology, Inc.; Watertown, MA) vertical runway. Two high-speed cameras (XC-1M; Xcitex Inc., Woburn, MA) were mounted to capture bird movement at 125 Hz from a lateral view and posterior view. Forces and video data were synchronized using the Procapture system (Xcitex Inc., Woburn, MA).

Following data collection, all SRFs were normalized using the animal’s body weight (% bw) to allow statistical comparison between individuals, and standardized for the direction of travel following Hanna and colleagues (5). A markerless pose estimation program DeepLabCut (15) was utilized to obtain positional data of three anatomical landmarks on the animal (7) and two points on the runway separated by a known distance. These positional data were used to calculate whole-body speed (m/s) and individual beak, hindlimb, and tail power (W kg^−1^; 3).

A series of ANCOVAs with whole-body speed as a covariate and Tukey’s post-hoc tests were used to assess statistical differences between the peak fore-aft and tangential force and average power of the hindlimb(s), beak, and tail. All variables were log-transformed prior to analysis. A complete description of these methods can be found in Supplement 1.

## Acknowledgments

The authors would like to thank Drs. Callum Ross and J.D Laurence-Chasen for helpful discussion during the initial stages of manuscript preparation. We also thank Christopher Hanna, Hannah Fischer, Elizabeth Davoli, and Allen Currier III for assistance with data collection. We thank Aaron Bastian for assistance with preparing the supplemental video. Finally, we thank animal husbandry staff. Without their help this study would not have been possible. This study was funded by the Center for Biomedical Innovation.

## Data Availability

All data used in this study is provided as a supplemental document.

