## Supplemental Methods for "Overcoming a “forbidden phenotype”: The parrot’s head supports, propels, and powers tripedal locomotion"

**This PDF file includes:**

Supplementary text
Legends for Movies S1
Legends for Datasets S1
SI References

**Other supplementary materials for this manuscript include the following:**

Movies S1
Datasets S1

**Supplementary Information Text**

### Supplementary Methods

#### Data collection

We collected substrate reaction force from six young adult rosy-faced lovebirds (*Agapornis roseicollis*; body mass range: 44.9 – 53.0 g; average: 48.8 ± 2.8 g) during vertical climbing. The target behavior was limited to vertical climbing as it is well-documented that psittaciforms utilize the beak during vertical ascent (1–5). Further, experiments altering substrate orientation reveal beak-use to be ubiquitous on vertical substrates, while the use of the beak is more intermittent at more acute angles (unpublished data). Animal protocols were approved by the New York Institute of Technology Institutional Animal Care and Use Committee (Protocol number: 2021-MG-01). Animals were free from any gait abnormalities and pathologies throughout the duration of study.

Beak, hindlimb, and tail fore-aft (Fx) and tangential forces (Fz) substrate reaction forces were collected (1250 Hz) as the birds climbed an instrumented vertical runway measuring 0.61 m in length and 0.15 m in width. The instrumented portion of the runway consisted of an Advanced Mechanical Technology, Inc. (AMTI) small-load force plate (model HE6X6; Watertown, MA) mounted with a custom three-dimensional printed platform (0.045 m length x 0.076 m width) to ensure recording of single beak, hindlimb, tail contact. The instrumented portion was flush mounted with the remainder of wooden runway and covered with shelf liners to improve gripping ability. Two high-speed cameras (XC-1M; Xcitex Inc., Woburn, MA) were mounted to capture bird movement at 125 Hz from a lateral view and posterior view. Forces and video data were synchronized using the Procapture system (Xcitex Inc., Woburn, MA).

We analyzed only the trials in which there was clear separation between forces generated by the beak, hindlimb(s) and tail. We refer to contact by any above-mentioned body part on the force platform as a “hit”. Trials in which more than one hit was registered were included as long as there was clear separation between the beak hindlimb(s) and tail contacts. All trials began by zeroing out all existing forces acting on the HE6X6 force plate.

**Data analysis and processing**

The HE6X6 force plate collects and outputs the force measurements in eight separate channels. Using a conversion formula provided by AMTI, a custom-written MATLAB (MathWorks, Natick, MA) code was used to export these data in the form of fore-aft (Fx) and tangential (Fz) forces across time. As standardization for the analysis of substrate reaction force during vertical climbing remains elusive, we elected to follow the terminology and data analysis practices of Hanna and colleagues (6). The animal moving upward on the vertical runway registered as a positive fore-aft force (+ Fx) whereas the animal applying force downward registered a negative fore-aft force (- Fx). The animal pushing into the force plate registered as a positive tangential force (+ Fz), while pulling away from the substrate registered as a negative tangential force (- Fz). Once standardized for the direction of travel, all substrate reaction forces were filtered through a low-pass Fourier filter at 15 Hz and normalized using the animal’s body weight (%bw) to allow statistical comparison between individuals (see below). Peak fore-aft and tangential forces were extracted from each beak, hindlimb, and tail hit.

A markerless pose estimation program DeepLabCut (7) was utilized to obtain positional data of the animals’ eye, front of wing, and tail tip in the lateral view throughout the extent of each trial. Anatomical landmarks were chosen based on Fujita and colleagues (8). DeepLabCut (7) uses a machine learning algorithm to accurately predict the positional data of points of interest and overcomes the need to label the points of interest in each trial video frame by frame. Approximately 150-200 frames of the bird's eye, front of wing, and tail tip as well as two additional points of a known distance were labeled to train the neural network. This known distance was used to calibrate the space and conversion factor used in subsequent velocity calculations.

Calculation of individual beak, hindlimb, and tail power (W) during locomotion followed protocols established by O'Connor and colleagues (9) and required the *x* and *y* positional data of the animals' eye, front of wing, and tail tip, the fore-aft force data, and information about the temporal boundaries of each hit. Using temporal boundaries, our custom-written MATLAB code output body part specific velocities (m/s) through time, average (whole-body) velocity (m/s) of the animal between temporal boundaries, power (W) through time, and average power (W) of each hit. Individual eye, front of wing, and tail tip velocities were used to calculate power for beak, hindlimb, and tail hits, respectively, as those landmarks, rather than whole-body velocity of the bird, more accurately reflected the velocity of the animal during each specific hit. Power was calculated by multiplying the average fore-aft force by the average velocity per hit and normalized with the animal's body weight (W $kg^{-1}$).

**Statistical Analysis**

All statistical analyses were conducted using R (10). For purposes of statistical analyses, the absolute value of the beak tangential forces we used. Shapiro-Wilk and Levene's tests were used to determine normality of data sets (11). All peak force data (Fx and Fz) and power underwent rank transformations prior to any statistical comparisons (11). As speed is well-known to influence the magnitude of substrate reaction force data (12, 13), least squares regressions were performed to determine whether peak forces and power were correlated to velocity. All variables were significantly affected by speed, and subsequent ANCOVAs were used to compare peak forces and power per hit.

**Movie S1 (separate file).** Tangential (top) and fore-aft (middle) forces and mechanical power (bottom) collected from the beak (orange), hindlimb (blue) and tail (green) of a climbing rosy-collared lovebirds (*Agapornis roseicollis*).

**Dataset S1 (separate file).** Peak fore-aft and tangential forces (%bw), average fore-aft velocity, and average mechanical power (W $kg^{-1}$) each for beak, hindlimb, and tail trial.
